# Characterization of Induced Susceptibility Effects of Soybean Aphid on Soybean

**DOI:** 10.1101/2020.07.25.221275

**Authors:** Surendra Neupane, Adam J. Varenhorst, Madhav P. Nepal

**Affiliations:** Department of Biology & Microbiology, South Dakota State University, Brookings, SD, USA; Department of Agronomy, Horticulture and Plant Science, South Dakota State University, Brookings, SD, USA 57007

**Keywords:** *Aphis glycines*, biotypes, RNA-seq, Soybean, Induced susceptibility

## Abstract

Soybean aphid (SBA) is one of the major pests of soybean (*Glycine max*) in the United States of America. The main objective of this research was to characterize interactions between two different biotypes of soybean aphids in susceptible and resistant soybean cultivars. Demographic and transcriptomic responses of susceptible and resistant (*Rag1*) soybean cultivars to aphid feeding were investigated in soybean plants colonized by aphids (biotype 1) in the presence or absence of inducer population (biotype 2) at day 1 and day 11. Leaves tissues collected at day 1 and day 11 post infestation were used for RNA sequencing, and ten RNA datasets with 266,535,654 sequence reads were analyzed. In the presence of inducer population, we found 746 and 243 differentially expressed genes (DEGs) in susceptible and resistant cultivars, respectively at day 1, whereas 981 and 377 DEGs were found in susceptible and resistant cultivars, respectively at day 11. Enrichment analysis showed a response to chitin, lignin catabolic and metabolic process, asparagine metabolic process, response to chemical unique to treatment with no inducer population, whereas, response to reactive oxygen species, photosynthesis, regulation of endopeptidase activity unique to treatment with inducer population. Furthermore, 14 DEGs were observed in *Rag* QTLs regions, particularly six DEGs in *Rag1* containing QTL. The identified DEGs in the experiment in both resistant and susceptible cultivars during the interaction of soybean and SBA are potential candidates for furthering investigation into induced susceptibility.

## INTRODUCTION

Invasive species have severely affected the agriculture system in numerous ways such as reducing yields and increasing costs of managing them affecting integrated pest management (IPM) [1,2]. *Aphis glycines* Matsumura (Hemiptera: Aphididae), the soybean aphid (SBA), a common invasive pest of soybean [*Glycine max* (L.) Merr] was first reported in North America in 2000 [3]. It is regarded as a common insect pest in China and many Asian countries [4]. In 2003, soybean aphid spread over 21 states of the U. S. and three Canadian provinces [5]. By the season of 2009, soybean aphid developed in the eastern region (New York and Ontario, Canada) beginning from July through August, as well as in Midwestern region (Minnesota, Wisconsin, and Iowa) spreading to 30 different states of the U.S. [6,7]. For an effective management approach, soybean lines that are naturally resistant to the aphids can be used to control SBA. Many researchers surveyed soybean germplasm collection and have identified soybean lines that have shown resistance to *A. glycines*. The resistance mechanism of the plant can be implemented in controlling pests without disturbing the environment [8]. Various dominant and recessive resistance to *A. glycines* (*Rag*) loci have been identified in soybean lines through various genetic analysis. Up to now, 16 *Rag* QTLs (reviewed in Neupane et al 2019 [9]) have been reported in various soybean plant introductions (PI).

Despite the identification of many monogenic and oligogenic genes for host plant resistance, the discovery of virulent biotypes of *A. glycines* that can survive on resistant varieties has been a serious threat. It has been estimated that the soybean cultivar with alone *Rag* and combined *Rag1* and *Rag2* can diminish the *A. glycines* growth by 34% and 49% respectively [10]. Up to now, four biotypes of aphid (biotype 1, biotype 2, biotype 3, and biotype 4) have been prevalent in the U.S having capability to reproduce in susceptible as well as resistant cultivars (with single or multiple *Rag* genes) [11–13]. Hence, the diverse population of both virulent and avirulent that appear phenotypically similar can engender induced susceptibility on the resistant plants [14]. The interaction between insect herbivores with their own host creates the condition called induced susceptibility that assists other consequent herbivores [15]. This type of susceptibility takes place between conspecifics on susceptible as well as resistant plants [15,16]. The phenotype of conspecific can be both virulent and avirulent biotype. Few studies have been done to understand induced susceptibility in *A. glycines* to answer the reason for a high number of *A. glycines* population in resistant soybean cultivars in North America. Varenhorst et al. 2015 [17] showed that virulent *A. glycines* increase the suitability of resistant soybean for avirulent conspecifics.

The comprehensive understanding of the transcriptomes would help in understanding the molecular interactions between soybean and *A. glycines*. A number of studies have been carried out using RNA-seq to unravel the molecular interactions for soybean-*A. glycines* herbivory with different objectives [18–20]. Brechenmacher et al., 2015 [20] identified 396 differentially regulated proteins and 2361 significantly regulated genes in different time response (up to 48 hours) soybean aphid infestation using two *Rag*2 and/or *Rag*2 near-isogenic lines of soybean. Among them, a gene of unknown function, a mitochondrial protease, and NBS-LRR gene those map within *Rag*2 locus are significantly upregulated in the presence of aphids. Prochaska et al., 2015 [18] identified 3 and 36 differentially expressed genes (DEGs) at day 5 and day 15 in resistant cultivar (KS4202), respectively whereas 0 and 11 DEGs at day 5 and day 15 in susceptible cultivar (K-03-4686), respectively. Most of the DEGs were related to WRKY transcription factors, peroxidases, and cytochrome p450s. Previously, Li et al. 2008 [21] studied soybean defense response to *A. glycines* generating transcript profiles using cDNA microarrays. In this study, they identified 140 genes related to the cell wall, transcription factors, signaling and secondary metabolism in response to resistance using resistant (cv. Dowling) and susceptible (cv. Williams 82) soybean cultivars. Studham and MacIntosh 2013[22] used oligonucleotide microarrays to study soybean-*A. glycines* interaction using aphid-resistant LD16060 with *Rag1* gene and aphid-susceptible SD01-76R. They identified 49 and 284 DEGs in 1 day of infestation (doi) and 7 doi in susceptible cultivar, respectively whereas 0 and 1 DEGs in 1 doi and 7 doi in resistant cultivar respectively studying transcript profiles determined after 1 and 7 days of aphid infestation. They suggested that the response of defense genes in the resistant plants are in constitutive in nature whereas, in susceptible plants, the defense genes are elicited only upon aphid infestation. This study is aimed to characterize induced susceptibility in soybean through the analysis of the transcriptional response of soybean in the presence of biotype 1 and biotype 2 soybean-aphids. Results of the study would have implication in soybean-aphid management and developing soybean cultivar with durable resistance to *A. glycines*.

## MATERIALS AND METHODS

### Plant Material and Aphid Colonies

Two genotypes of soybean were used: susceptible soybean cultivar was LD12-15838R and the resistant cultivar was LD12-15813Ra. The resistant cultivar contains *Rag1* QTL. These genotypes were infested with two aphid populations defined namely biotype 1 (avirulent) and biotype 2 (virulent biotype [11]). The biotypes are defined by the response to *Rag1* genes and were identified in Illinois [11]. The biotype 1 and biotype 2 populations originated from a colony maintained by Iowa State University (Ames, IA). Among them, colonies of biotype 1 originated from field populations in Ohio and were maintained in a colony at the Ohio State University biotype. At South Dakota State University, aphid colonies were maintained using susceptible cultivar SD01-76R for biotype 1 and resistant cultivar LD12-15813Ra for biotype 2. The aphid populations used in this study were randomly selected removing the leaves from the soybean plants used for maintaining the colonies.

### Greenhouse Experiment

To characterize induced susceptibility effects, randomized complete block design (RCBD) greenhouse experiment was conducted using twelve treatments, three replications (plants) in three blocks (nine experimental units per treatment). We followed the treatments as explained by the procedure by Varenhorst et al 2015[17]. The initial feeding population of *A. glycines* was termed as an inducer population and the subsequent feeding population of *A. glycines* was termed as a response population. Three seeds of LD12-15838R and LD12-15813Ra were planted into damp soil (Professional Growing Mix, Sun Gro Horticulture, MA, USA) in each pot of dimension of 10.1 cm by 8.89 cm (500 ml; Belden Plastics, MN, USA). Pots were placed onto plastic flats (87 × 15 × 5 cm). The soybean plants were watered filling the flats when top soil began to dry. The plants were thinned down to one plant per pot upon reaching the V1 developmental growth stage. V2 staged soybean plants (Day 0) were infested with avirulent inducer populations using with a combination of zero inducer [23], 50 *A. glycines* (50 avirulent), or 50 *A. glycines* (50 virulent) onto a ventral side of a middle leaf of first trifoliate except the control plants. The infested trifoliate was covered with a small no-see-um mesh net (Quest Outfitters, Sarasota, FL) and secured with the paper clip and tangle trap to confine within the first trifoliate of the plants. After 24 hrs. (Day 1), one-day leaves from second trifoliate were collected from one replication set of each block and snap frozen in the liquid nitrogen. After sample collection from one replication, response population of 15 *A. glycines* (15 avirulent), or 15 *A. glycines* (50 virulent) were added upon the middle leaf of second trifoliate (except on sampled and control plants). The whole plants were covered with the large no-see-um mesh net (Quest Outfitters, Sarasota, FL) to confine movement of aphids between the plants. The response population was allowed to move freely about the plant with the exception of first trifoliate. This ensures the spatial isolation of inducer and response populations. The response populations were counted on each plant to confirm the colonization by the response populations on day 5. On day 11, the response population of aphids was counted and the day 11 leaf samples from the one replication sets of each block were collected and snap frozen in the liquid nitrogen. The samples were kept at −80°C for further analysis. The greenhouse conditions were maintained approximately 24-25°C and a 16-hour photo period (16 light: 8 dark).

The aphid counts (response population) collected at 11^th^ day after the inducer infestations were analyzed using R statistical software version 3.2.4 (https://www.r-project.org/). The main effects of the inducer population, soybean cultivar, and the interaction of inducer population by soybean cultivar were analyzed using the model Response Counts ~ Inducer + Inducer: Cultivar. We checked the effect of both treatment and block for susceptible and resistant cultivars separately. The model Aphid Counts ~ Treatment + Block was applied in the analysis of variance [24]. The treatment means based on *A. glycines* numbers were separated using Fisher- least significant difference (LSD) test at *P* < 0.05 using agricolae package [25] in R. The average SBA counts were plotted using GraphPad Prism 8.0.2 (San Diego, California USA, www.graphpad.com).

### RNA Extraction, Library Construction, and RNA-sequencing

RNA was extracted from the leave samples from resistant and susceptible cultivars treated with no aphids, biotype 2: biotype1 collected at day 1 and no aphids, biotype 2: biotype1 and no aphids: biotype1 at day 11. Briefly, leaf samples from each treatment were grounded in liquid nitrogen with pestle and mortar to a fine powder followed by their processing for total RNA extraction using PureLink RNA mini kit (Invitrogen, USA). RNA samples were treated with TURBO™ DNase (Invitrogen, USA) to remove any DNA contamination following the manufacturer’s instructions. Assessment of the isolated RNA integrity was performed by 1% agarose gel electrophoresis, and RNA concentration was measured by Nanodrop 2000 (Thermo Fisher Scientific, USA). Three replicates from these treatments in resistant and susceptible cultivars were pooled in equimolar concentration. The cDNA libraries were constructed and sequenced at South Dakota State University Sequencing Facilities. RNAseq library construction was prepared using Illumina’s TruSeq Stranded mRNA Kit v1 (San Diego, CA). The libraries were quantified by QuBit dsDNA HS Assay (Life Technologies, Carlsbad, CA). The libraries were sequenced on an Illumina NextSeq 500 using a NextSeq 500/550 High Output Reagent Cartridge v2 (San Diego, CA) at 75 cycles.

### RNA-seq Analysis

Quality control of reads was assessed using FastQC program (version 0.11.3) (https://www.bioinformatics.babraham.ac.uk/projects/fastqc/) [26]. Low quality bases (QC value < 20) and adapters were removed by trimming using the program Trimmomatic (version 0.36) [27] (options: PE -phred33 LEADING:3 TRAILING:3 SLIDINGWINDOW:4:15 HEADCROP:8 MINLEN:30). High-quality single-end reads were mapped against the primary coding sequences of *G. max*. The coding sequences (*Gmax:Gmax_275_Wm82.a2.v1.transcrip t_primaryTranscriptOnly.fa.gz*) were obtained from the Phytozome database and aligned using Salmon v0.9.1 [28]. The read quants were filtered with 0.5 counts per million (CPM) in at least one sample. The quantified raw reads were transformed using regularized log (rlog) which is implemented in the DESeq2 package. The transformed data were subjected to exploratory data analysis such as hierarchical clustering, K-means clustering, principal component analysis (PCA), and visualization of clusters using the t-SNE map. Gene co-expression networks were constructed for divided datasets with the weighted gene co-expression network analysis (WGCNA) package [29] using following parameters: most variable genes to include-3000 genes, soft threshold- 4, minimum module size- 20. The quantified transcript reads obtained from Salmon were employed in CLC Genomics Workbench 9.5 to obtain the differentially expressed genes (DEGs) using Karl’s z-test with false discovery rate (FDR) <0.01 and log2fold change more than a 2-fold. The annotations of the DEGs were obtained from Soybase [30] (www.soybase.org). To understand the molecular pathways enriched GO Biological processes, GO Cellular, GO molecular function, and KEGG pathways for DEGs were analyzed using a graphical enrichment tool REVIGO [31], ShinyGO [32] and integrated Differential Expression and Pathway analysis (iDEP 0.81, R/Bioconductor packages) [33]. The enriched transcription factor binding motifs in promoters in different comparisons were identified in 300bp upstream of DEGs using both iDEP and ShinyGO [32]. The biological relevance of DEGs was visualized using MapMan [34]. The total transcripts of soybean were first converted to bins using the Mercator tool [35] and uploaded to MapMan to assign bins to each differentially expressed transcript.

## RESULTS

### Effect of Inducer Populations on the Response Populations

The hypothesis of response population being positively affected by the presence of inducer population or conspecifics was tested considering the main effects of the inducer population, soybean cultivar, and the interaction of inducer population by soybean cultivar. The response population was significantly affected by the main effects of inducer population (*F* = 15.821, df = 1, *P* = 0.000130) and cultivar (*F* = 11.642, df = 1, *P* = 0.000926). Induced susceptibility effect on both susceptible and resistant soybean cultivars, as we observed increased response population densities in the 50 virulent inducer population treatments compared to the none inducer population treatment (**Figure 1a** and **1b**). Also, the interaction of inducer population on soybean cultivar was significant (*F* = 3.956, df = 1, *P* = 0.049386) as the response population in the resistant cultivar was lower than that of a susceptible cultivar.

**Figure 1.**
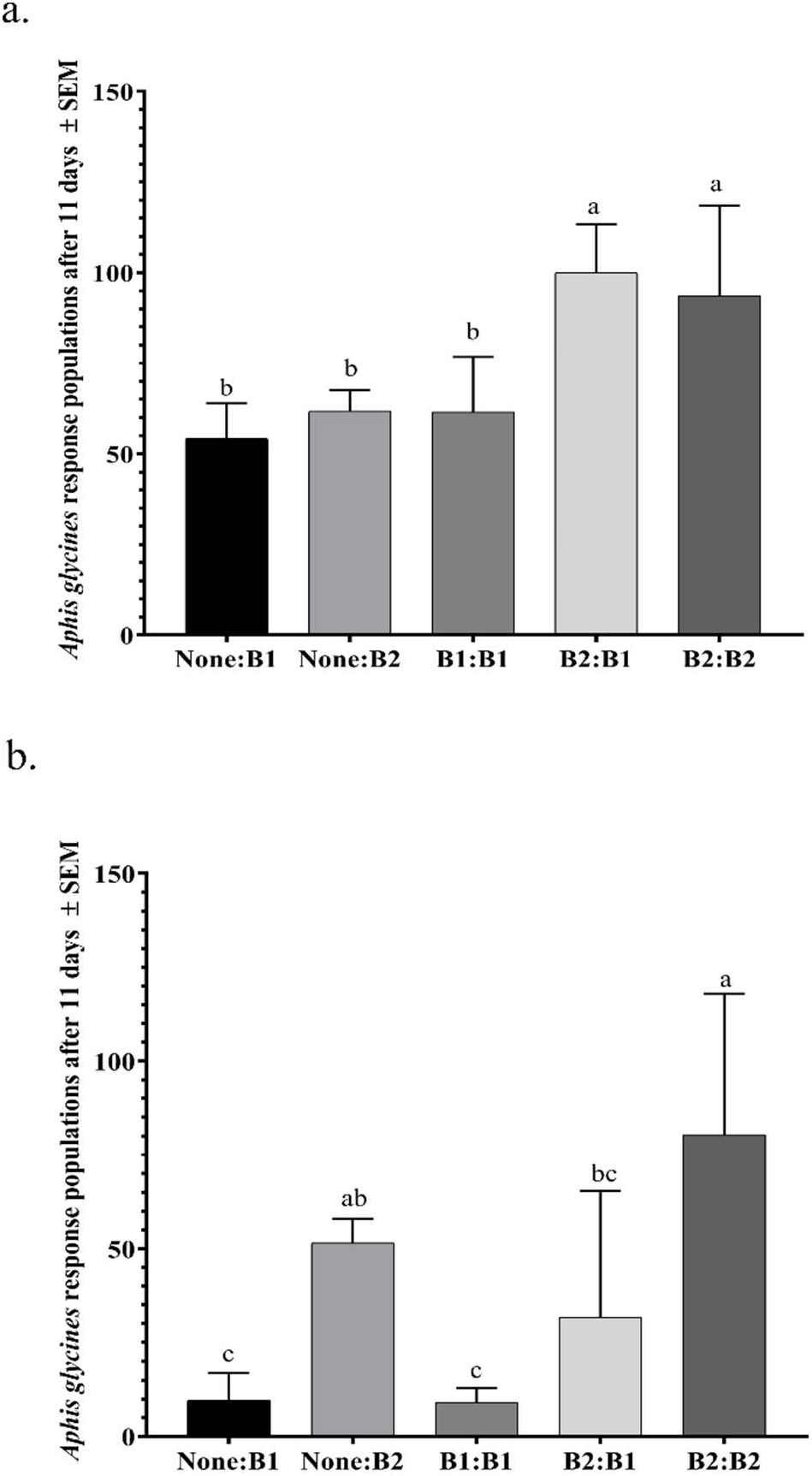
Effect of avirulent (B1) and virulent (B2) inducer populations on avirulent (B1) and virulent (B2) response populations on both (a) susceptible and (b) resistant soybean. For this experiment, the susceptible soybean cultivar was LD12-15838R and the resistant cultivar was LD12-15813Ra. Lowercase letters indicate significance among treatments (*P* < 0.05). Plotted values represent the means of the avirulent response population.

Upon application of model Response counts ~ Treatment + Block was applied in analysis of variance [24], both treatment (*F* = 10.950, df = 5, *P* = 6.92e-07) and block (F= 4.497, df = 2, *P* = 0.0167) effect were significant in susceptible cultivars. Whereas, block effect was insignificant (*F* = 0.588, df = 2, *P* = 0.56) in resistant cultivars. Thus, we applied a reduced model Response counts ~ Treatment in resistant cultivars. One way ANOVA was applied to observe the significance of treatment (*F* = 7.601, df = 5, *P* = 2.52e-05) in resistant cultivars. Fisher-least significant difference (LSD) test at *P* < 0.05 was applied to see the separation of treatment means based on *A. glycines* numbers. In susceptible cultivars, we observed the separation of means of avirulent (response) population between the treatments with none, biotype 2 as an inducer with biotype 1 as an inducer. Response populations for the biotype 2 as inducer population treatments were 84.4% greater than the response population that received the “none inducer” treatment in the susceptible cultivar. In resistant cultivars, we did not observe the separation of means of avirulent (response) population between the treatments with zero, biotype 2 as an inducer with biotype 1 as an inducer. However, response populations for the biotype 2 as inducer population treatments were 228% greater than the response population that received the “none inducer” treatment in the resistant cultivar.

### RNA-seq Analysis

A total of 10 RNA libraries were prepared and sequenced with the sequencing depth ranging from 24,779,816 to 29,72,4913. Total reads of 266,535,654 were subjected to FastQC analysis to determine the data quality. The Phred quality scores per base for all the samples were higher than 30. Upon mapping these reads, we obtained high mapping rate ranging from 90.4% to 92.9%. Among the mapped reads, 85.8% to 91.9% reads were uniquely mapped.

### WGCNA Analysis

The co-expression networks were used to detect correlated networks of genes and their enrichment in the divided datasets to compare difference on the day 1 and day 11 treatments. Weighted gene co-expression network analysis identified a network of 3,000 genes divided into 11 co-expression modules in four day 1 samples, and a network of 2,999 genes divided into 15 co-expression modules in six day 11 samples (**Figure 2**, **Supplementary File**[36]). In entire modules, the enrichment analysis found several highly enriched KEGG pathways for day 1 and day 11 samples. The only KEGG pathways enriched in day 1 samples, but not in day 11 samples were Phenylpropanoid biosynthesis, Glycine, serine and threonine metabolism, Alpha-Linolenic acid metabolism, Glycolysis / Gluconeogenesis, and Cysteine and methionine metabolism. Whereas, the only KEGG pathways enriched in day 11 samples, but not in day 1 samples were Plant-pathogen Interaction, Flavonoid biosynthesis, MAPK signaling pathway, Glucosinolate biosynthesis, and Alanine, aspartate and glutamate metabolism, Cutin, suberine, and wax biosynthesis (**Table S1[36]**). The common pathways for both time points included Biosynthesis of secondary metabolites, Metabolic pathways, Ribosome, Porphyrin, and chlorophyll metabolism.

**Figure 2.**
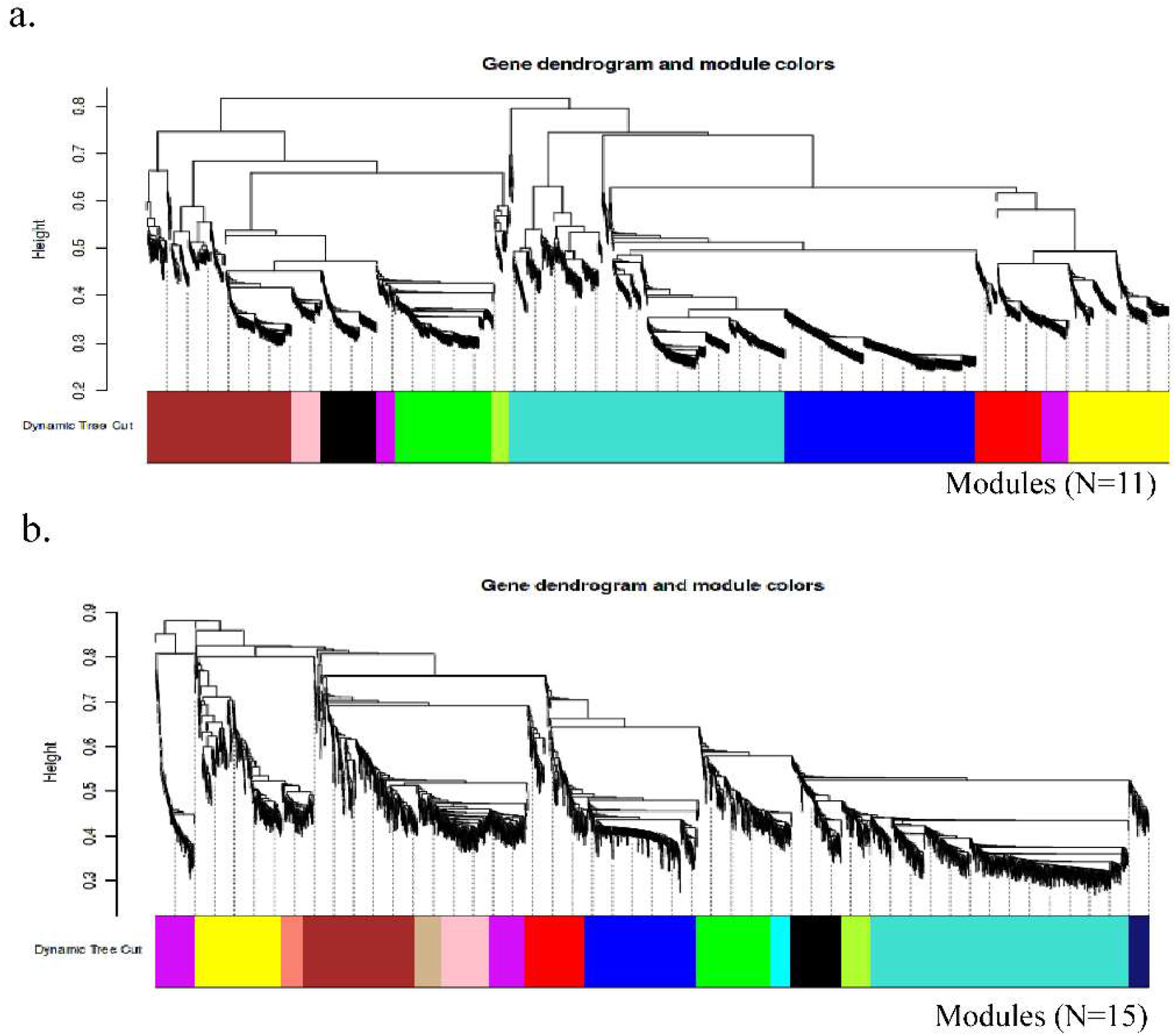
Weighted gene co-expression network analysis identified a network of 3,000 genes divided into 10 co-expression modules in day 1 samples (a), and a network of 2,999 genes divided into 15 co-expression modules in day 11 samples (b).

### Hierarchical and K-means Clustering

After filtering with 0.5 counts per million (CPM) in at least one sample and rlog transformation, a total of 37,468 genes (66.9% of original 55,983) were retained for clustering and visualization. We used gene clustering to assess if day 11 had more gene clusters enriched for defense-related pathways than day 1 samples. K-means clustering identified five clusters of correlated genes in day 11 samples (**Figure 3**), and Cluster A was enriched primarily with the biosynthesis of secondary metabolites, Cluster B was enriched with various plant defense-related pathways such as MAPK signaling pathway, Plant-pathogen interaction, and plant hormone signal transduction (Supplemental File [36]). Four clusters were identified (A-D) in day 1 samples, of which Cluster A was enriched with photosynthesis, carbohydrate metabolism, Cluster B was enriched with fatty acid metabolism, glucosinolate biosynthesis, Cluster C and D were enriched with biosynthesis of secondary metabolites, and plant defense-related pathways such as MAPK signaling pathway (Supplemental File[36]). Promoter analysis of clusters in day 1 samples found 80 enriched transcription factor binding motifs in four clusters (A, B, C, and D). Enriched binding motifs consisted of twelve transcription factor families: AP2, AT hook, bHLH, bZIP, CG-1, CxC, Homeodomain, Myb/SANT, NAC/NAM, TBP, TCP, and WRKY. Promoter analysis in day 11 samples found 100 enriched transcription factor binding motifs, consisting of eight transcription factor families: AP2, bHLH, bZIP, CG-1, E2F, LOB, Myb/SANT, and TCP (Supplemental File 2). Six transcription factor families (AP2, bHLH, bZIP, CG-1, Myb/SANT, TCP) were found in both time periods. Four transcription factor families were unique to day 1 samples (AT hook, CxC, Homeodomain, and NAC/NAM), whereas, two transcription factor families were unique to day 11samples (E2F and LOB).

**Figure 3.**
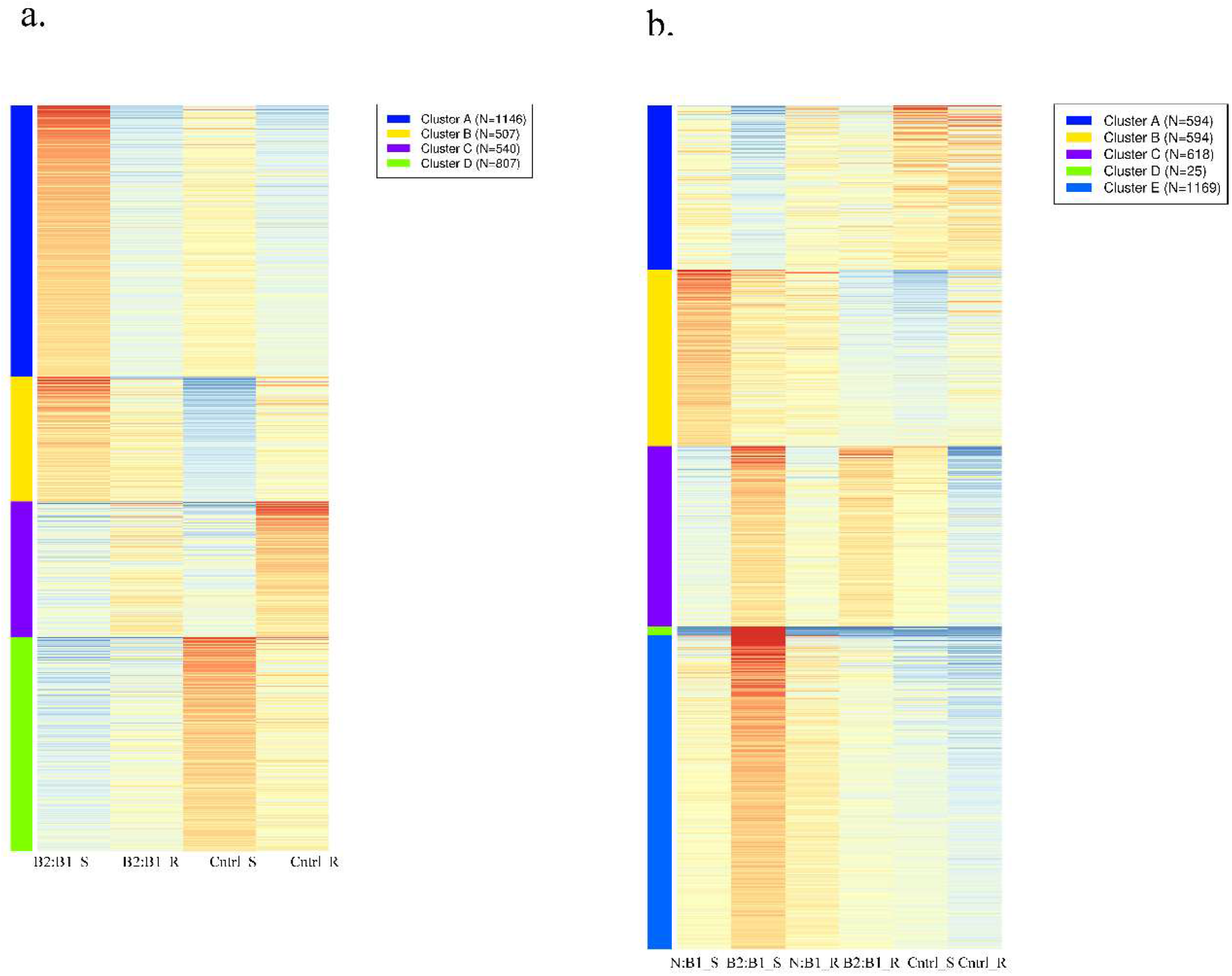
K-means clustering of top 3,000 most highly variable genes in Day-1 samples (a) and Day-11 samples (b).

### Gene Expression Analysis

The pair wise comparisons between treatments in two different treatments (none: B1 and B2: B1) at day 1 and day 11 with FDR < 0.01 and log2fold-change > 2 as cutoffs resulted differentially expressed genes (DEGs) shown in **Table 1**. We further investigated these genes using Venn diagrams (**Figure 4**). At day 1, we found 746 and 243 DEGs in susceptible and resistant cultivars, respectively treated with biotype 2 as inducer and biotype 1 as response population (B2: B1). Whereas 981 and 407 DEGs were found in susceptible and resistant cultivars, respectively at day 11 treated with biotype 2 as inducer and biotype 1 as response population (B2: B1). At day 11 we found 520 and 377 DEGs in susceptible and resistant cultivars treated with no inducer and biotype 1 as a response population (none: B1). In total, at day 11, we found 1,274 and 638 DEGs in susceptible and resistant cultivars, respectively upon comparing treatments with none: B1 and B2: B1.

**Table 1.** A number of up-regulated and down regulated DEGs in different comparisons.

| Time | Comparisons | Cultivar | Up regulated | Down regulated |
| --- | --- | --- | --- | --- |
| Day 1 | B2:B1 vs Control | Susceptible | 364 | 382 |
|  | B2:B1 vs Control | Resistant | 239 | 4 |
| Day 11 | none:B1 vs Control | Susceptible | 196 | 324 |
|  | B2:B1 vs Control | Susceptible | 660 | 321 |
|  | none:B1 vs Control | Resistant | 154 | 223 |
|  | B2:B1 vs Control | Resistant | 214 | 196 |

**Figure 4.**
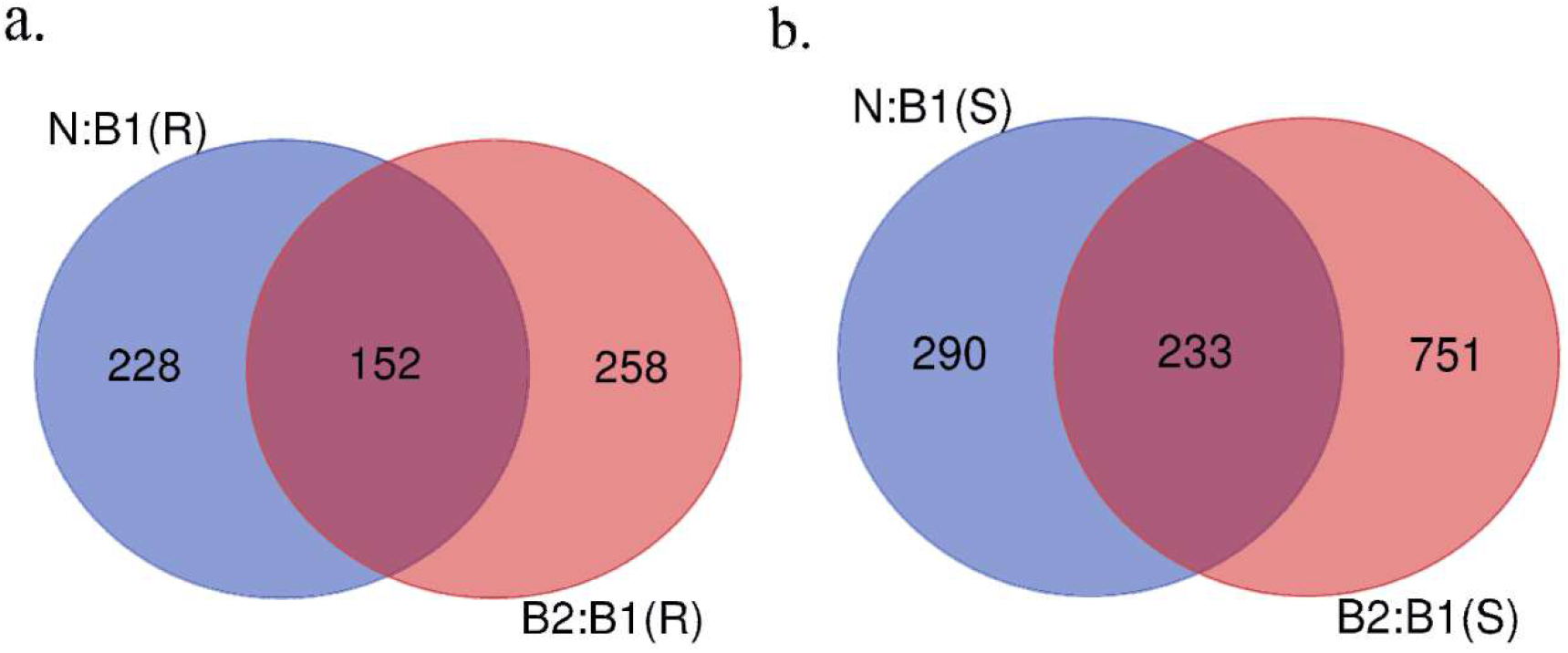
Venn diagram showing DEGs for two treatments N: B1 (none: B1) and B2: B1 in resistant (a) and susceptible cultivars (b).

### GO, KEGG Enrichment and MapMan Analysis

The 1,274 and 638 DEGs in susceptible and resistant cultivars, respectively were subjected to GO enrichment analysis for biological process, molecular function, and KEGG pathways. In susceptible cultivar, the DEGs were enriched for various biological processes including Jasmonic acid mediated signaling pathway, Response to chitin, Phenylpropanoid metabolic process, Regulation of defense response, Response to chemical or organic substance, Response to wounding, Hormone metabolic process, Reactive oxygen species metabolic process, Regulation of macromolecule biosynthetic and metabolic processes. Among them, Jasmonic acid mediated signaling pathway was unique to none: B1 treatment and Phenylpropanoid biosynthetic process and Glucosinolate metabolic process were unique to B2: B1 treatment (**Figure 5**). In terms of KEGG pathways, these genes were enriched for Cutin, suberine and wax biosynthesis (FDR=5.36E-07), Biosynthesis of secondary metabolites (FDR=5.36E-07), Glucosinolate biosynthesis (FDR=1.04E-05), Phenylpropanoid biosynthesis (FDR=4.89E-05), MAPK signaling pathway (FDR=8.84E-05), Plant hormone signal transduction (FDR=0.047596525) and others represented in **Figure S1[36]** and Supplementary File[36]. Promoter analysis of 1,274 DEGs showed 30 enriched transcription factor binding motifs. Enriched binding motifs consisted of seven transcription factor families: AP2, bHLH, bZIP, CG-1, LOB, SBP, and TCP (Supplemental File 3).

**Figure 5.**
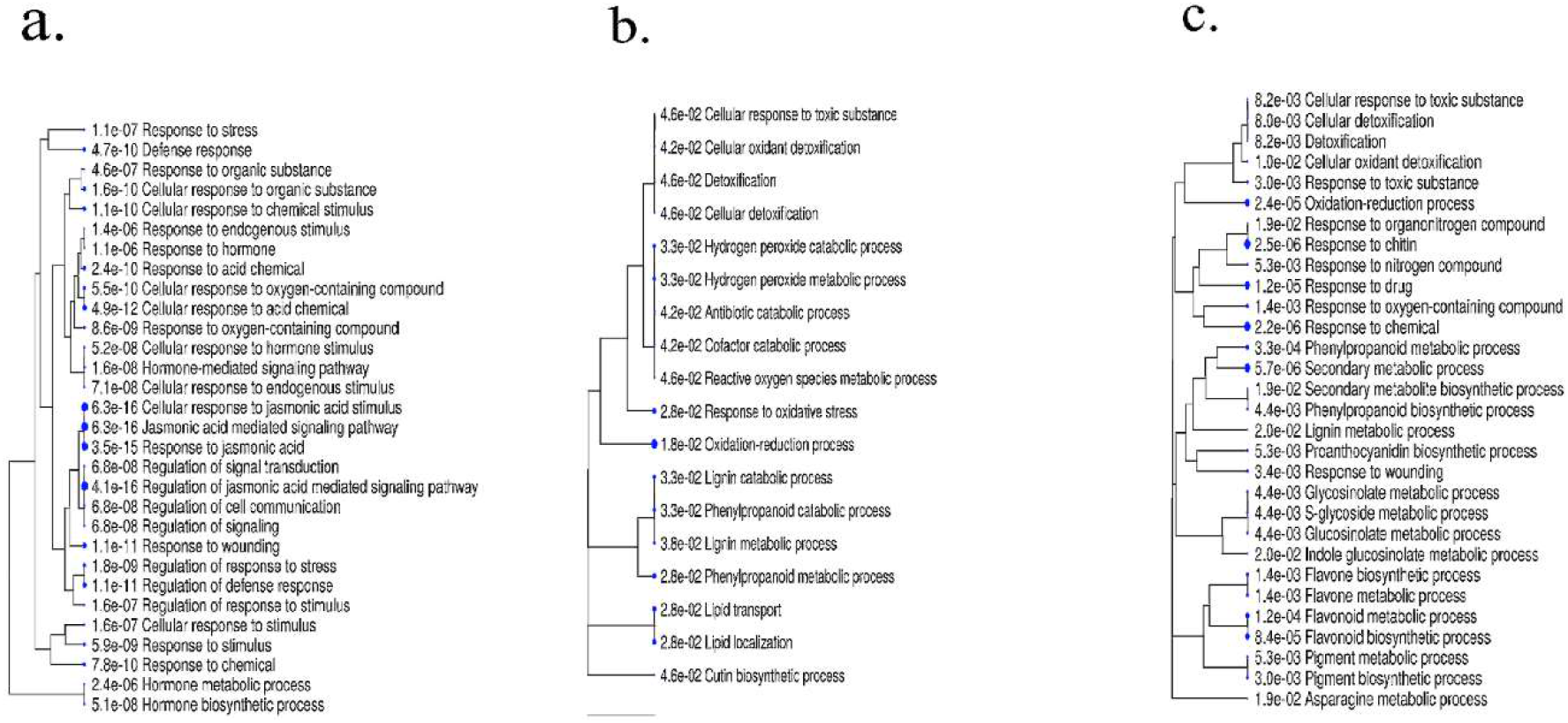
Enriched GO biological processes specific to treatments in susceptible cultivar at day 11. None: B1 (a), common (b), and B2: B1(c).

Differentially expressed genes visualized using biotic stress pathway integrated into MapMan showed distinct expression patterns in none: B1 and B2: B1 treatments in both susceptible and resistant cultivars. The biotic stress overview pathway generated by MapMan demonstrated the involvement of multifaceted defense related genes in the presence of inducer and no inducer population in both susceptible and resistant plants. In susceptible reaction, 280 (of 523) with 26 bins and 362 (of 984) DEGs with 25 bins were associated with the biotic stress pathway for none: B1 and B2: B1 treatments, respectively (**Figure 6**). As compared to treatment none: B1, upregulated genes related to abiotic stress (bin 20.2), peroxidases (bin 26.12), abscisic acid hormone pathway (bin 17.1), respiratory burst (bin 20.1.1), glutathione S transferase (bin 26.9), pathogenesis related (PR)-proteins (bin 20.1.7), and secondary metabolism (bin 16), and heat shock proteins (HSPs) (bin 20.2.1).

**Figure 6.**
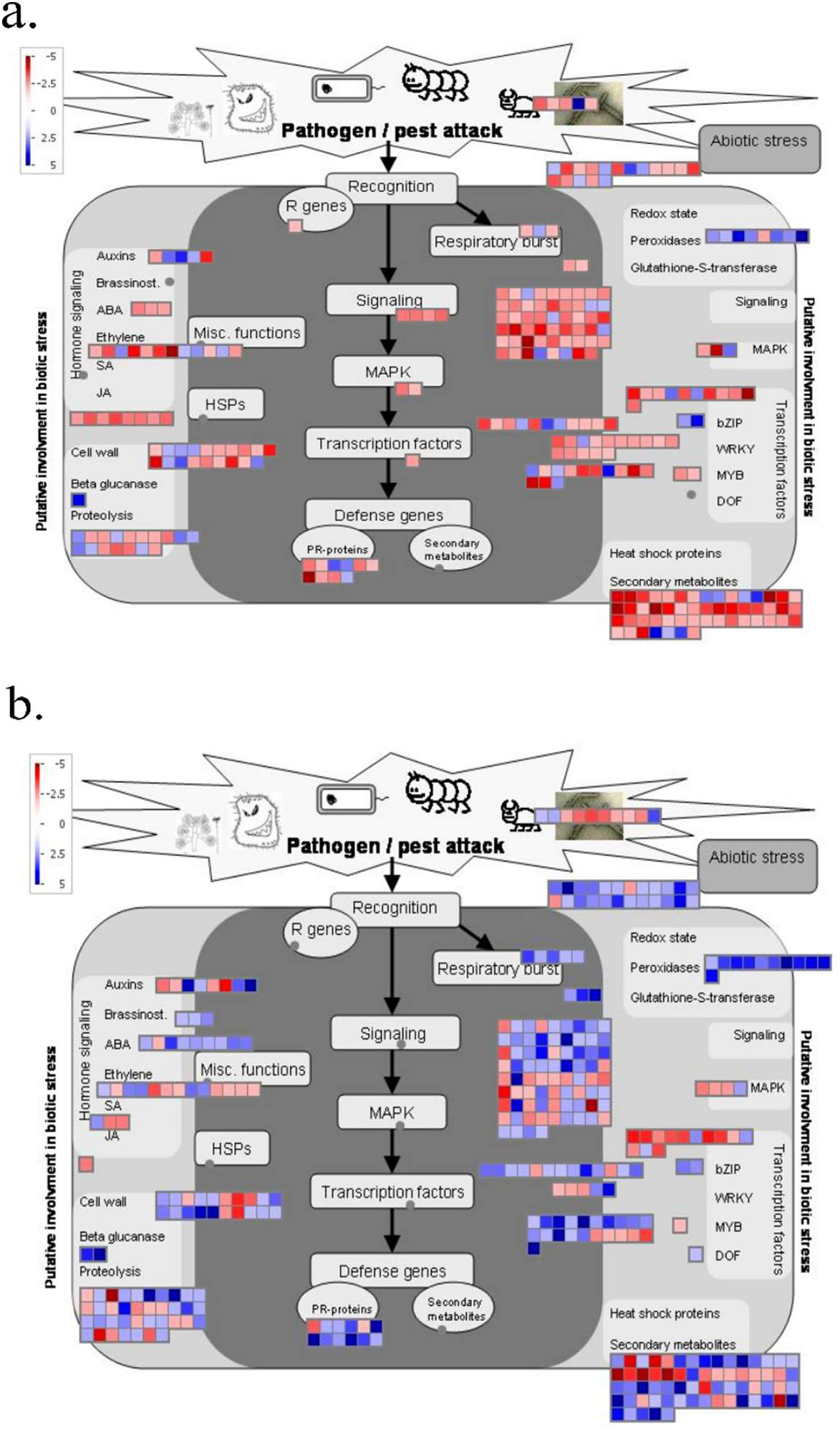
Biotic stress pathway overview of differentially expressed genes in susceptible cultivar at day 11. None: B1 (a) and B2: B1 (b). Blue color indicates the up-regulated and red color indicates the down regulated genes. False discovery rate (FDR) *p* < 0.01 and log2fold change ≥ 2 or ≤ −2 were used to identify the differentially expressed genes.

Whereas, 638 DEGs in resistant cultivar were particularly enriched for Photosynthesis, Lignin metabolic process, negative regulation of endopeptidase activity, response to cytokinin, Inositol catabolic process which were different from the susceptible reaction (**Figure S_2[36]_**). Among them, response to chitin, lignin catabolic and metabolic process, asparagine metabolic process, response to chemical were unique to none: B1 treatment, whereas, response to reactive oxygen species, photosynthesis, and regulation of endopeptidase activity were unique to B2: B1 treatment (**Figure 7**). These genes were enriched for KEGG pathways such as Photosynthesis (FDR= 0.005883), Glutathione metabolism (FDR=0.009895), Cutin, suberine and wax biosynthesis (FDR=0.012764), Cysteine and methionine metabolism (FDR=0.046797), and Flavonoid biosynthesis (FDR=0.046797) (Supplementary File[36]). Promoter analysis of 638 DEGs showed 30 enriched transcription factor binding motifs. Enriched binding motifs consisted of four transcription factor families: bHLH, bZIP, CG-1, and TCP (Supplemental File[36]).

**Figure 7.**
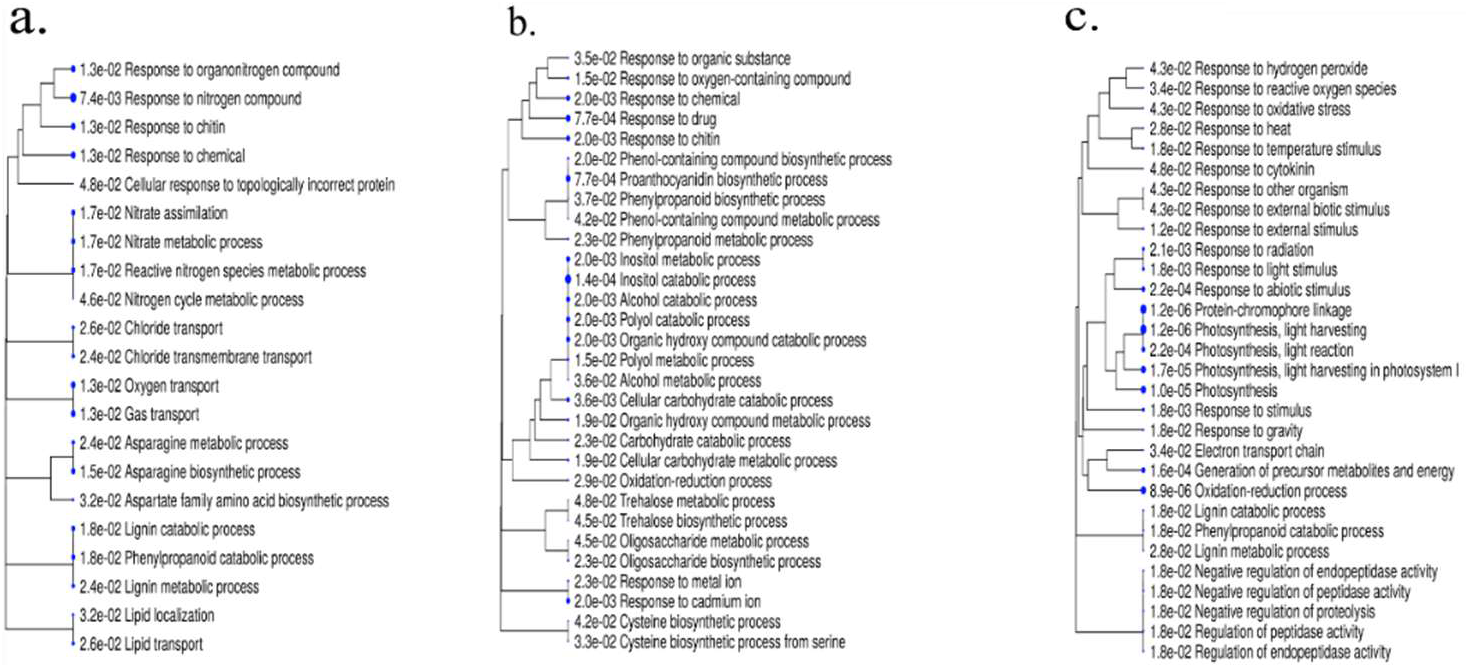
Enriched GO biological processes specific to treatments in resistant cultivar at day 11. None: B1 (a), common (b), and B2: B1 (c).

In resistant reaction, MapMan biostress pathway revealed 154 (of 380) with 21 bins and 176 (of 410) DEGs with 23 bins associated with the biotic stress pathway for none: B1 and B2: B1 treatments, respectively (**Figure 8**). As compared to treatment none: B1, upregulated genes related to transcription factors [WRKY (bin 27.3.32), MYB (27.3.25)], peroxidases (bin 26.12), abscisic acid hormone pathway (bin 17.1), respiratory burst (bin 20.1.1), glutathione S transferase (bin 26.9), salicylic acid hormone pathway (bin 17.8), jasmonic acid hormone pathway (bin 17.7), pathogenesis related (PR)-proteins (bin 20.1.7), and secondary metabolism (bin 16), and heat shock proteins (HSPs) (bin 20.2.1).

**Figure 8.**
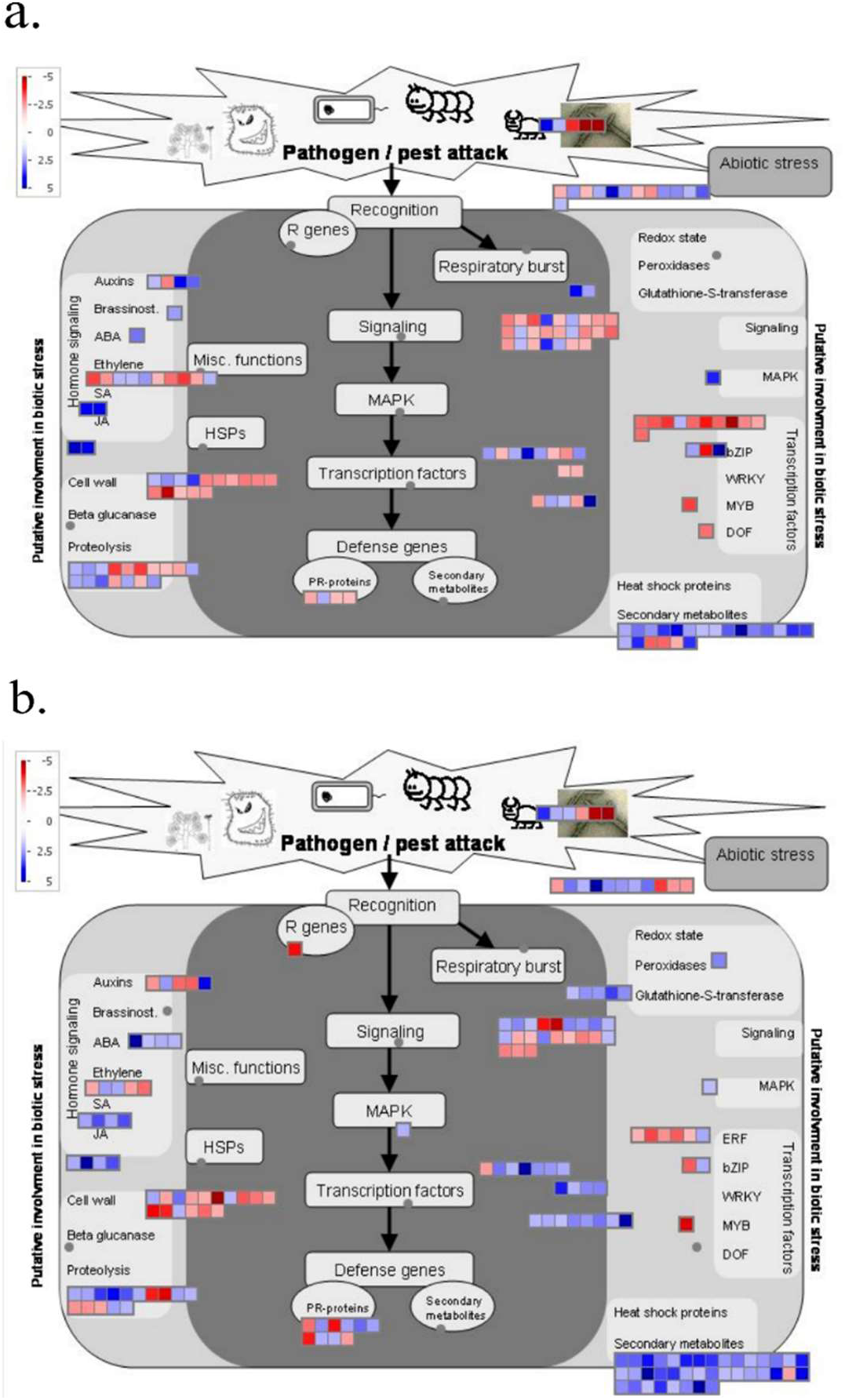
Biotic stress pathway overview of differentially expressed genes in resistant cultivar at day 11. None: B1 (a) and B2:B1 (b). Blue color indicates the up-regulated and red color indicates the down regulated genes. False discovery rate (FDR) *p* < 0.01 and log2fold change ≥ 2 or ≤ −2 were used to identify the differentially expressed genes.

### Comparison of the DEGs between Two-time Points

Further, we compared DEGs for samples treated with biotype 2 as inducer and biotype 1 as response population (B2: B1) on day1 and day 11. Of 626 DEGs in resistant cultivar, 216 were unique to day 1 samples, 383 were unique to day 11 samples and 27 were expressed at both time points (**Figure 9a**). Likewise, of 1,621 DEGs in susceptible cultivar, 637 were unique to day 1 samples, 872 were unique to day 11 samples and 112 were expressed at both time points (**Figure 9b**).

**Figure 9.**
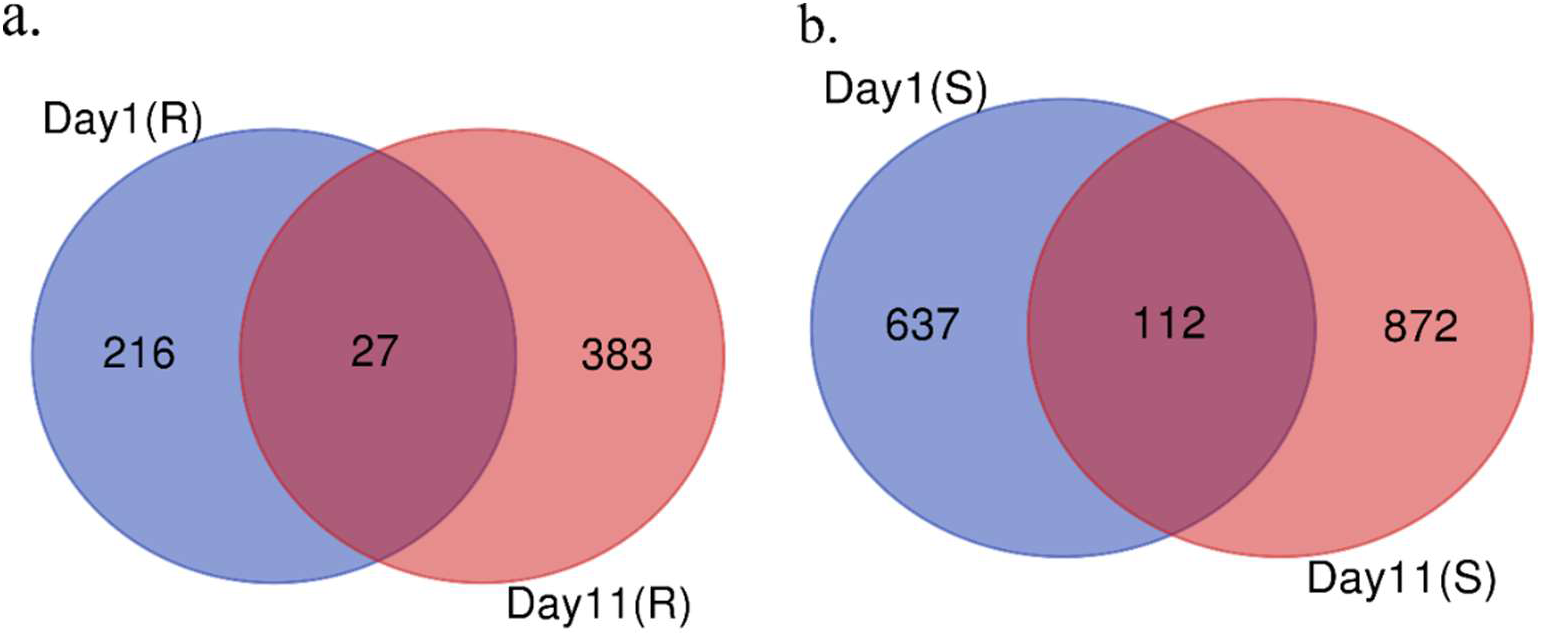
Venn diagram showing DEGs for treatment B2: B1 at day 1 and day 11 in resistant (a) and susceptible (b) cultivars.

At day 1, MapMan biostress pathway revealed 284 (of 749) with 24 bins and 90 (of 243) DEGs with 16 bins associated with the biotic stress pathway in susceptible and resistant cultivars with B2: B1 treatment, respectively (**Figure 10**). As compared to a susceptible reaction, the resistant reaction showed fewer bins associated with the biostress pathway with almost all upregulated genes.

**Figure 10.**
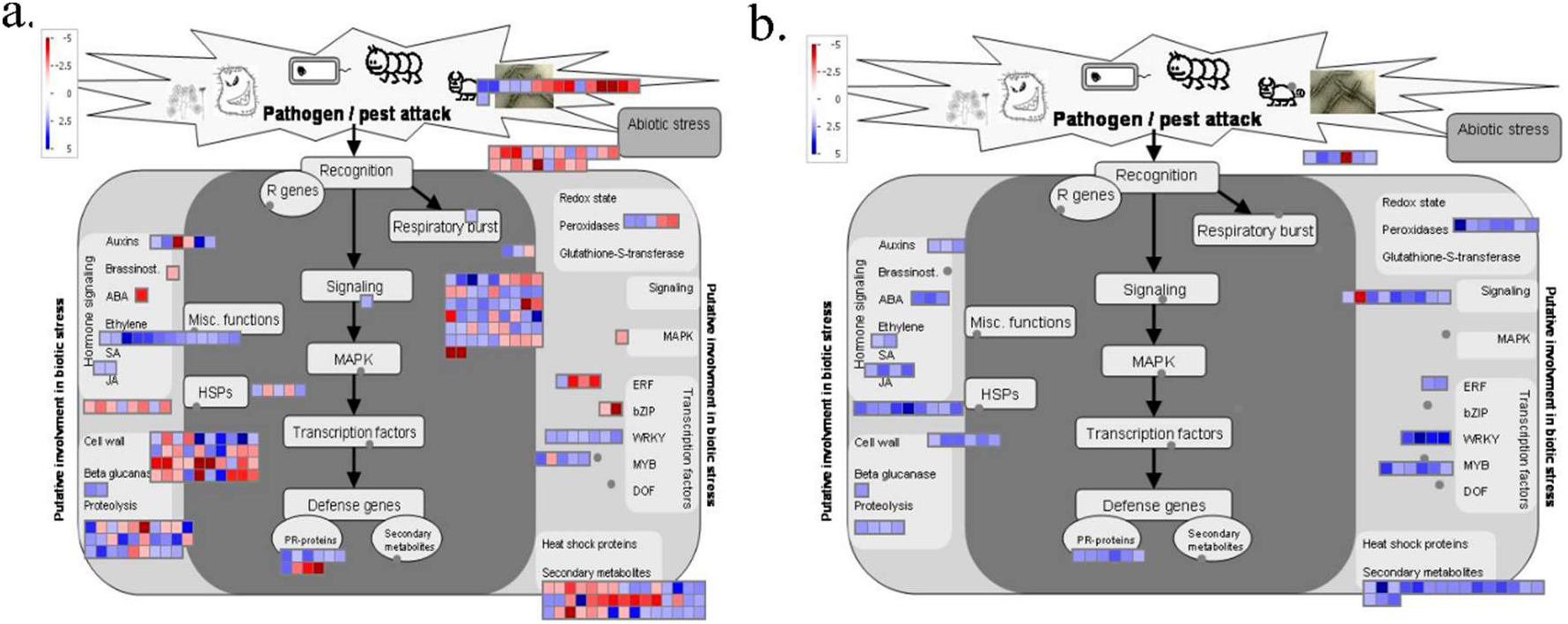
Biotic stress pathway overview of differentially expressed genes at day 1 with B2: B1 treatment. Susceptible (a) and resistant (b). Blue color indicates the up-regulated and red color indicates the down regulated genes. False discovery rate (FDR) *p* < 0.01 and log2fold change ≥ 2 or ≤ −2 were used to identify the differentially expressed genes.

The 27 overlapped DEGs in resistant cultivar represented the variation in the level of expression at day 1 and day 11 (**Table 2**). In both cultivars, we observed no difference, increased and decreased expression pattern of the genes (**Table 2** and **Table S2[36]**). Particularly in resistant cultivar, elicitor-activated gene 3-2 (Glyma.01G021000), jasmonic acid carboxyl methyltransferase (Glyma.02G054200), Calcium-binding EF-hand family protein (Glyma.02G108700), 2-oxoglutarate (2OG) and Fe(II)-dependent oxygenase superfamily protein (Glyma.18G273200) showed static level or no difference of expression at day 1 and day 11. Whereas, the expression of NAD(P)-binding Rossmann-fold superfamily protein (Glyma.03G222600) increased from 2.74 to 5.48 log2fc, myo-inositol oxygenase 1 (Glyma.08G199300) from 2.22 to 3.42 log2fc, Aluminium induced protein with YGL and LRDR motifs (Glyma.15G072400) from 2.05 to 4.26 log2fc, basic helix-loop-helix (bHLH) DNA-binding superfamily protein from 2.61 to 4.68 log2fc, and ribonuclease 1 (Glyma.20G036100) from 3.70 to 7.00 log2fc. In contrast, the expression of drought-repressed 4 (Glyma.03G068200) was decreased from 2.46 to −6.07 log2fc, nitrate reductase 1 (Glyma.13G084000) from 2.11 to −5.01, pathogenesis-related 4 (Glyma.19G245400) from 2.42 to −3.05 log2fc. At day 1, particularly, seven genes belonging to peroxidases and six cytochrome P450s were highly upregulated. In addition, disease resistance-responsive (dirigent-like protein) family protein (Glyma.03G045600, Glyma.08G019900, Glyma.12G030300) were expressed in the range of 2.6 to 3.3 log2foldchange, Kunitz family trypsin and protease inhibitor protein (Glyma.08G235300, Glyma.16G212500, Glyma.08G235400) were expressed in the range of 2.2 to 4.7 log2foldchange, laccase 3 (Glyma.02G231600, Glyma.14G198900) were expressed in the range of 2.8 to 3.1, TRICHOME BIREFRINGENCE-LIKE 27 (Glyma.01G034600, Glyma.02G031400) and TRICHOME BIREFRINGENCE-LIKE 42 (Glyma.12G233500) were expressed in the range of 2.0 to 2.1 log2foldchange, Ferritin/ribonucleotide reductase-like family protein (Glyma.16G056300) was expressed by 3.129 log2foldchange, WRKY24 (Glyma.18G238600) was expressed by 4.2 log2foldchange (Supplementary File[36]). At day 11, particularly, wall associated kinase 5 (Glyma.13G035900) was expressed by 2.78 log2fold change, glutathione S-transferases (Glyma.18G043700, Glyma.11G212900, Glyma.07G139600, Glyma.07G139900, Glyma.07G139700) were expressed in the range of 2.0 to 3.5 log2foldchange, Toll interleukin-1 receptor-like Nucleotide-binding site Leucine-rich repeat (TNL) genes (Glyma.03G048600, Glyma.03G052800, Glyma.03G048700, Glyma.03G047700) expressed in the range of 2.4 to 3.1 log2foldchange, senescence-related genes (Glyma.06G273600, Glyma.13G222100, Glyma.15G090100, Glyma.16G052000) were expressed in the range of 2.1 to 4.1 log2foldchange, UDP-glucosyltransferases (Glyma.08G109100, Glyma.02G105000, Glyma.10G062600, Glyma.20G196000) were expressed in the range of 2.1 to 3.65 log2foldchange, myo-inositol oxygenases (Glyma.07G013900, Glyma.05G224500) were expressed in range of 3.5 to 3.9 log2foldchange, ferritin 4 (Glyma.02G262500) was expressed by 2.26 log2foldchange, WRKY40 (Glyma.17G222300) and WRKY67 (Glyma.03G002300) were expressed in the range of 2.1 to 2.7 log2foldchange (Supplementary File[36]).

**Table 2.**
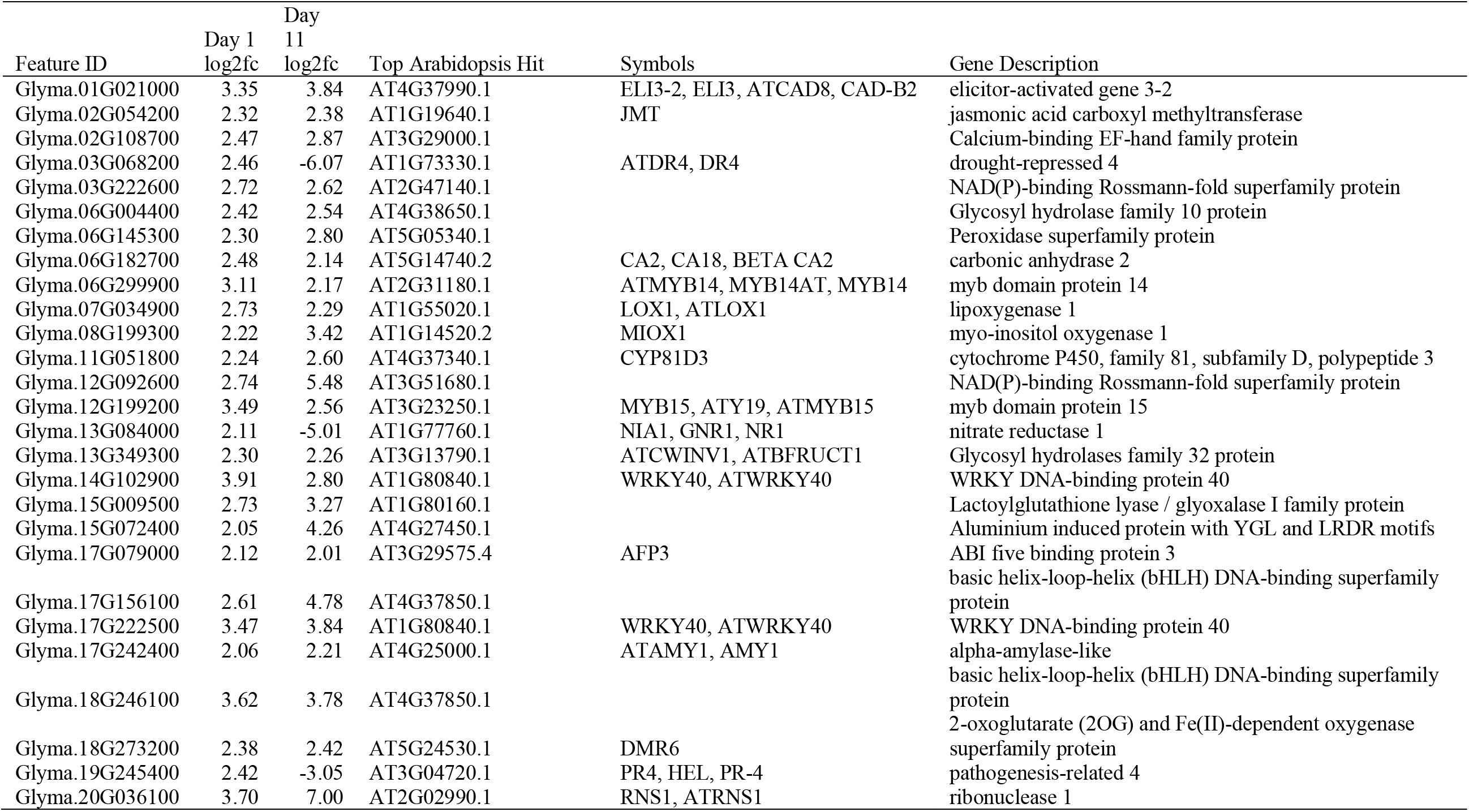
List of 27 common DEGs for treatment B2:B1 at day 1 and day 11 in a resistant cultivar.

### DEGs Coincident with Rag QTL Genes

We identified 1,691 non-redundant genes in the *Rag* QTLs and compared with the DEGs found in the resistant cultivar, LD12-15813Ra. We found 14 DEGs that were coincident with the *Rag* QTL genes with lipoxygenase 1 (Glyma.07G034900) being up-regulated at both day 1 (2.73 log2foldchange) and day 11 (2.29 log2foldchnage) treated with B2 as an inducer population and B1 as a response population and Gibberellin-regulated family protein (Glyma.17G237100) downregulated at day 11 in both treatment conditions none: B1 and B2: B1. Protein kinase family proteins with leucine-rich repeat domain (Glyma.16G065600) and Gibberellin-regulated family protein (Glyma.17G237100) were downregulated at day 1 and day 11, respectively treated with B2 as an inducer population and B1 as a response population. Likewise, arabinogalactan protein 22 (Glyma.07G087200) and Gibberellin-regulated family protein (Glyma.17G237100) were downregulated treated with no inducer population and B1 as a response population at day 11 (**Table 3**).

**Table 3.**
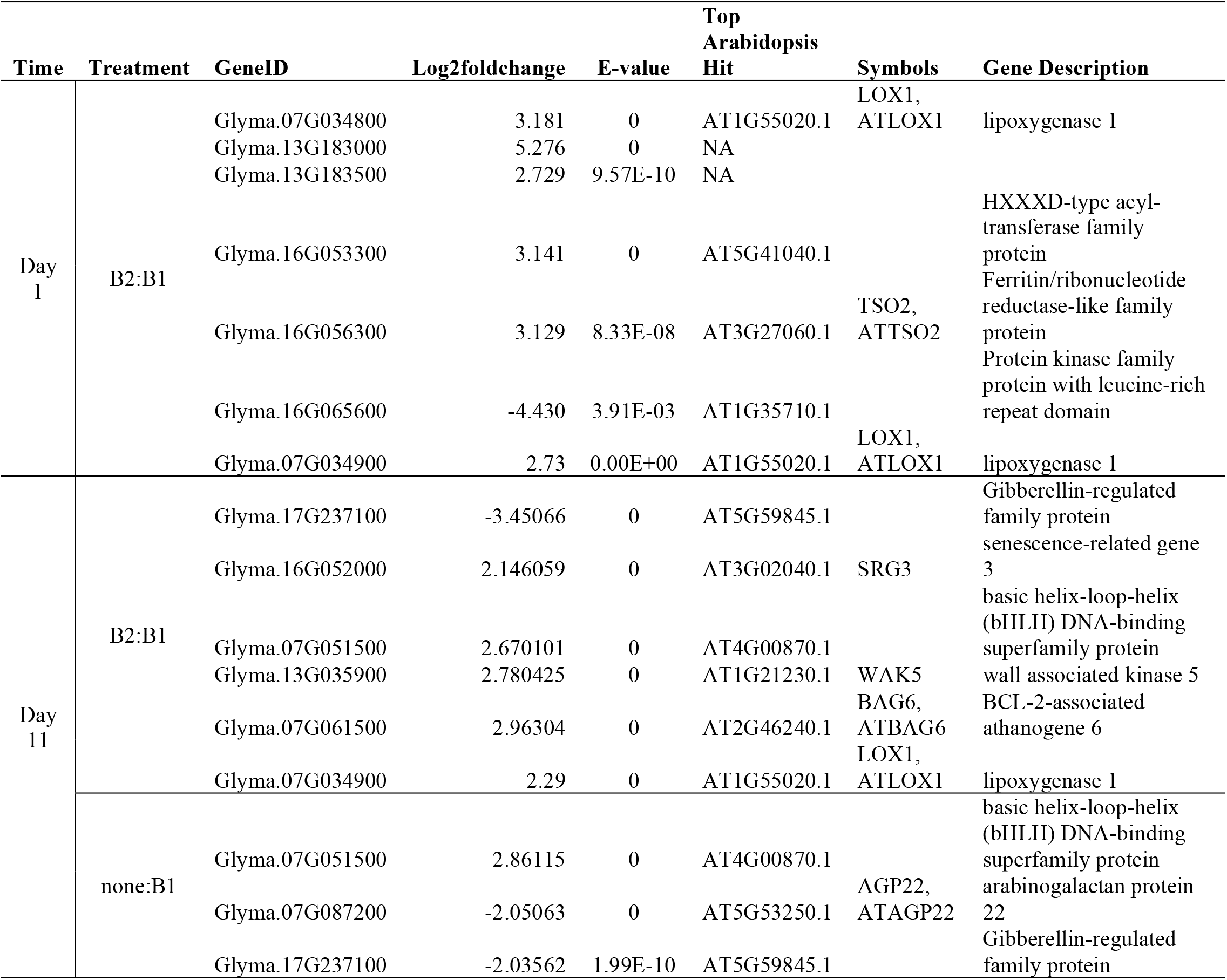
List of DEGs coincident with *Rag* QTL genes

## DISCUSSION

This experiment is the first attempt to characterize the induced susceptibility effect that promotes the avirulent *A. glycines* populations in both resistant and susceptible cultivar treated with virulent inducer populations. Previously, this effect was initially tested with *Rag1* + *Rag*2 (IA3027RA12) cultivar and subsequent tests in near-isogenic soybean cultivars containing no *Rag* genes (IA3027), *Rag1* (IA3027RA1), using biotype 1 and biotype 2 soybean aphids in a growth chamber and semi-field settings [17]. We first validated this effect using susceptible soybean cultivar (LD12-15838R) with no *Rag* gene and the resistant cultivar (LD12-15813Ra) with *Rag1* gene in the greenhouse settings with slight modifications on response population density (15 instead of five). In the meantime, we collected leaves samples for the transcriptomic study. We observed both ‘feeding facilitation’ [37] and ‘obviation of resistance’ [16] which are the two subcategories of induced susceptibility. Feeding facilitation refers to the condition where conspecifics are favored on either susceptible or resistant host plants in presence of herbivore, irrespective of its genotype. Whereas obviation of resistance refers to the condition where avirulent conspecifics on the resistant plant are favored in the resistant host plant in the presence of virulent herbivore. We chose treatments with no aphids (control), biotype 2: biotype1 (B2: B1) and no aphids: biotype1 (none: B1) collected at day 1 and day 11 for the transcriptomic study. These treatments were chosen as we expect some insights on gene expression pattern in resistant and susceptible cultivars in time course response in presence or absence of virulent soybean aphids as an inducer population and avirulent soybean aphids as a response population. The day 1 samples were selected expecting some response to the host by the inducer population. The day 11 samples were selected as we expected both physical and metabolic changes caused by both inducer and response populations.

The initial WGCNA analysis revealed 11 and 15 co-expression modules on day 1 and day 11, respectively enriched for various pathways in both resistant and susceptible cultivars. At day 1 or 24 hours, we found an enriched pathway for cysteine and methionine metabolism which was also enriched in the DEGs in resistant cultivar discussed below. Many plant species utilize *S*-methylmethionine and glutathione to transport sulphur molecules in the phloem [38]. Aphids might have an efficient mechanism for the production of methionine and cysteine from the phloem metabolites [39]. It has been shown that peach aphid and pea aphid in symbiosis with the endosymbiont, *Buchnera aphidicola* incorporate sulphur from inorganic sulphate transported to the phloem sap [39,40]. The presence of aphid endosymbiotic bacteria [41] in the aphids might be one of the causes for feeding facilitation and obviation of resistance by soybean aphids. The possibility of the role of endosymbionts including plant viruses and aphid effector molecules causing induced susceptibility was discussed by Varenhorst et al. 2015 [17]. Other enriched pathways at day 1 were Phenylpropanoid biosynthesis, Alpha-Linolenic acid metabolism, Fatty acid biosynthesis. Previous studies have shown that the phenylpropanoid pathway was induced in the resistant (*Rag1*) Dowling cultivar at 6 and 12 h after aphid feeding [42]. The pathways related to α-linolenic acid metabolism and fatty acid biosynthesis corresponds to the precursor for jasmonic acid pathway via the oxylipin pathway [43]. This shows that soybean aphids can induce hormone response inducing changes in fatty acid metabolism within 24 hours. The production of various phytohormones such as JA including SA and ET upon aphid infestation on the response of resistant (*Rag1*) and susceptible near-isogenic soybean lines [22]. Such effect was also seen in two soybean varieties (DK 27–52 and DK 28–52) when infested with soybean aphid in the field environment [44]. At day 11, we found an enriched pathway for Flavonoid biosynthesis, Plant pathogen interaction, MAPK signaling pathway, and Glucosinolate biosynthesis. The interaction of plant and pathogen involves pathogen-associated molecular patterns (PAMPs) of pathogens by pattern recognition receptors (PRRS) of the host [45]. These plant-parasite interactions have caused a battle in the molecular avenue where evolutionary arms race takes place [46]. There are various models that describe plant-pathogen interactions such as the gene for gene model, guard model, decoy model, bait and switch model, and zig-zag model [47–50]. Zig-zag model depicts the interaction between plant and parasites [48]. It is still unknown if aphid and other insects interaction follow the particular model [46]. As reviewed by Wu and Baldwin, 2010 [51] early defense signaling events take place in a cell of insect attacked leaf. Briefly, aphid elicitors are perceived by the receptors on plasma membrane trigger Ca^2+^ channels and produce Ca^2+^. Ca^2+^ binds with NADPH oxidase which gets enhanced through phosphorylation by CDPKs eventually producing reactive oxygen species (ROS). MAPK pathways are activated quickly among which SIPK and WIPK trigger the synthesis of Jasmonic acid (JA) and JA-Ile (JA-isoleucine) which is a central regulator of plant innate immunity [52]. Another enriched pathway at day 11 was Glucosinolate biosynthesis. The involvement of secondary metabolites such as glucosinolates have been documented in two separate studies as a defensive compound when *Myzus persicae* infested *Arabidopsis* for three [53] and seven days [54].

The K-means clustering revealed five and four clusters for day 11 and day 1 samples. The pathway enrichment analysis of the clusters supported the enrichment of entire modules obtained from WGCNA analysis. Enriched binding motifs of these clusters revealed AT hook [55], CxC [56], Homeodomain [57], and NAC/NAM [58] transcription factor families unique to day 1 samples. whereas, two transcription factor families were unique to day 11samples (E2F [59], and LOB [60]). Six transcription factor families (AP2 [61], bHLH [62], bZIP [63], CG-1[64], Myb/SANT [65], TCP [66]) were found in both time periods. Among them, AP2, bHLH, bZIP, CG-1, LOB, SBP, and TCP were particularly enriched in susceptible reaction whereas, bHLH, bZIP, CG-1, and TCP were enriched in resistant reaction.

At day 11, upon analyzing DEGs between treatments none: B1 and B2: B1 we observed a significant number of DEGs in susceptible cultivar than resistant cultivar (1,274 vs 638). The DEGs in susceptible cultivar were enriched for many biological processes related to defense programs such as MAPK signaling pathway, Plant hormone signal transduction, and Plant-pathogen interaction. These are the major components of the PTI program of defense mechanism. Such an effect in which significant induction of defense programs in susceptible cultivar aphid-susceptible SD01-76R when infested with soybean aphid for 21 days [67]. The DEGs in resistant cultivar were enriched for Photosynthesis, Glutathione metabolism, Cutin, suberine and wax biosynthesis, Cysteine and methionine metabolism, and Flavonoid biosynthesis. Particularly, glutathione metabolism was enriched in which one gene, glutathione peroxidase 6 (Glyma.01G219400) was upregulated by 2.12 log2foldchange in the none: B1 treatments. Whereas three genes belonging to glutathione S-transferases (Glyma.07G139700, Glyma.07G139900, Glyma.14G067200) were upregulated by 2.04 to 2.5 log2foldchange in the B2: B1 treatments. The structural damage on the host upon aphid feeding may be linked to the accumulation of excessive reactive oxygen species (ROS) in the attacked organs [68]. Plant glutathione S-transferases (GSTs) make such endogenous substrates and xenobiotics (e.g., ROS) less toxic upon adding glutathione molecule via nucleophilic or addition reactions [69]. We observed glutathione peroxidase6 being upregulated in none: B1 treatments. Sometimes, GSTs exhibit glutathione-peroxidase activity for the reduction of hydroperoxides [70]. The enrichment of Cysteine and methionine metabolism which was also observed in initial WGCNA analysis at day 1 was observed in DEGs in resistant cultivar at day 11. This shows that cysteine and methionine metabolism pathway is active from initial aphid feeding to the 11^th^ day. Another enriched pathway in the resistant cultivar was photosynthesis. Previous transcriptomic study on soybean near-isogenic lines differing in alleles for an aphid resistance gene, *Rag5* following infestation by soybean aphid biotype 2 has shown DEGs enriched for photosynthesis [71]. Physiologically, photosynthesis rates have been reduced up to 50% on soybean aphid infested leaflets [72].

The comparison of DEGs was further expanded to see a pattern of the expression of DEGs especially focusing to common and unique genes at day 1 and day 11 in the resistant cultivar when treated with biotype 2 as an inducer population. Particularly on day 1, we observed upregulation TRICHOME BIREFRINGENCE-LIKE 27 and −42 proteins and laccase 3. TRICHOME BIREFRINGENCE-LIKE genes contribute to the synthesis and deposition of the secondary wall [73]. Likewise, laccase genes also play a role in cell wall lignification [74]. The cell wall modification and deposition of callose are considered as the chemical defense responses during PAMP-triggered immunity (PTI) response after recognition of components from the aphid saliva [75]. The role of laccase in tolerance to the insect pests cotton bollworm (*Helicoverpa armigera*) and cotton aphid (*Aphis gosypii*) has been shown in cotton [76]. Upregulation of other genes at day 1 involved peroxidases, cytochrome P450s. The role of peroxidases in scavenging ROS during the defense mechanism has been clearly documented in plant-aphid interactions [77,78] including soybean-soybean aphid interaction [18]. Plant cytochrome P450s are importantly involved in jasmonic acid mediated plant defense in response to wound and insect attack [79]. Other DEGs belonged to disease resistance-responsive (dirigent-like protein) family protein, Kunitz family trypsin, protease inhibitor protein, and Ferritin/ribonucleotide reductase-like family protein. Previously, DIR-like proteins were also upregulated during feeding of spruce (*Picea* spp.) stem-by boring insects (i.e., white pine weevil, *Pissodes strobi*) in bark tissue and defoliating insects (i.e., western spruce budworm, *Choristoneura occidentalis*) in green apical shoots [80]. Kunitz family trypsin and protease inhibitor protein target various proteases of phytophagous pests and pathogen as a resistance response [81]. Previously, Kunitz family trypsin and protease inhibitor genes were reported as a differentially expressed gene in tolerant soybean cultivar upon soybean aphid feeding [18]. Another gene that encodes Ferritin/ribonucleotide reductase-like family protein was upregulated at day 1 response. The differential expression of ferritin as a resistance response has been shown in previous studies as a part of constitutive resistance mechanism in soybean-soybean aphid interactions [18,21,67]. Upregulation of ferritins in resistant plants can limit the availability of iron to the insect [67]. At day 11, four TNL genes, four homologs of WRKY40, one homolog of WRKY67, four senescence-related genes, four UDP-glucosyltransferases, two myo-inositol oxygenases, five glutathione S-transferases were uniquely upregulated in the resistant cultivar when treated with biotype 2 as an inducer. The expression of four TNL genes at day 11 shows the involvement of canonical resistance genes. Numerous plant resistance genes involved in defense mechanism encode proteins containing nucleotide-binding site (NBS) and a leucine-rich repeat (LRR) motifs [82]. For example, *Vat* gene, which confers resistance to *Aphis gossypii* in melon (*Cucumis melo*) is also an NBS-LRR gene [83].

We examined DEGs that are coincident with the 1,691 genes that were assessed from the *Rag* QTLs. The cultivar used in this experiment is LD12-15813Ra with *Rag1* gene. The mapping and inheritance mechanism of the *Rag1* gene has been well studied in various soybean cultivars [42,84–86]. *Rag1* loci were finely mapped as a 115 kb interval on chromosome 7 through genetic mapping using cultivar Dowling (PI 548663; donor parent of *Rag1*) and Dwight (PI587386; aphid-susceptible parent) [87]. We found 14 DEGs that were coincident with the *Rag* QTL genes. Among them, six genes belonged to *Rag1* QTLs. These genes belonged to lipoxygenase 1 (Glyma.07G034800, Glyma.07G034900) basic helix-loop-helix (Glyma.07G051500, Glyma.07G051500), BCL-2-associated athanogene (Glyma.07G061500) were upregulated while arabinogalactan protein 22 (Glyma.07G087200) was downregulated.

In summary, demographic datasets obtained from the greenhouse experiments validated induced susceptibility effects, which was further characterized by the genetic datasets obtained from RNA-seq technique. The characterization, however, was limited to two treatments: one with no inducer population and biotype 1 as a response population and another with biotype 2 as an inducer population and biotype 1 as a response population in both resistant and susceptible cultivars. Many DEGs were common and unique in two cultivars and treatments that were enriched for various biological processes and pathways and were functionally related to known defense mechanisms reported in various host-aphid systems. The responses to aphid biotype 1 infestation in presence or absence of inducer population at day1 and 11 revealed significant differences on the gene enrichment and regulation in resistant and susceptible cultivars. The assessment of DEGs in *Rag* genes QTLs, particularly in *Rag1* containing QTL on chromosome 7, six non-NBS-LRR genes – Glyma.07G034800, Glyma.07G034900, Glyma.07G051500, Glyma.07G051500, Glyma.07G061500, Glyma.07G087200 revealed distinct expression in treatments with absence or presence of inducer population at day 1 and day 11. However, four TNL genes – Glyma.03G048600, Glyma.03G052800, Glyma.03G048700, Glyma.03G047700 were upregulated in resistant cultivar treated with biotype 2 as an inducer population and biotype 1 as response population at day 11 which suggest their crucial role in the interaction effects. Further experiments based upon metabolomics, proteomics, and validation of the candidate genes will be needed to understand the mechanism underlying induced susceptibility effects.

## ACKNOWLEDGMENTS

Authors would like to acknowledge Philip Rozeboom and Alyssa Vachino for their assistance in greenhouse experiments. Funding for the greenhouse experiments and RNA sequencing came partly from South Dakota Soybean Research and Promotion Council (SDSRPC-SA1800238), and partly from the United States Department of Agriculture hatch projects (SD00H469-13 and SD00H659-18) to M. Nepal.

## DATA DEPOSITION

The data described in this study can be assessed from Neupane et al 2019 [88].

## CONTRIBUTIONS

SN carried out the experiments. AJV and MPN conceived the project and contributed designing the experiments. All authors contributed writing this manuscript.

